# Electrical Energy Storage with Engineered Biological Systems

**DOI:** 10.1101/595231

**Authors:** Farshid Salimijazi, Erika Parra, Buz Barstow

## Abstract

The availability of renewable energy technologies is increasing dramatically across the globe thanks to their growing maturity. However, large scale electrical energy storage and retrieval will almost certainly be a required in order to raise the penetration of renewable sources into the grid. No present energy storage technology has the perfect combination of high power and energy density, low financial and environmental cost, lack of site restrictions, long cycle and calendar lifespan, easy materials availability, and fast response time. Engineered electroactive microbes could address many of the limitations of current energy storage technologies by enabling rewired carbon fixation, a process that spatially separates reactions that are normally carried out together in a photosynthetic cell and replaces the least efficient with non-biological equivalents. If successful, this could allow storage of renewable electricity through electrochemical or enzymatic fixation of carbon dioxide and subsequent storage as carbon-based energy storage molecules including hydrocarbon and non-volatile polymers at high efficiency. In this article we compile performance data on biological and non-biological component choices for rewired carbon fixation systems and identify pressing research and engineering challenges.

## Background

The penetration of renewable electricity sources like wind, solar, and wave is significantly increasing across the world thanks to their growing maturity and an increasing pressure to control climate change. These same forces are also driving the electrification of transportation, considerably increasing demands on the electrical grid. However, it’s well known that unlike traditional electricity sources, the power output of most renewables is variable at best, and completely unreliable at worst [1]. In order to replace a large fraction of the current electricity supply with renewable sources and enable electrified transportation, electrical energy storage at low-cost and large scale will be essential.

How much electricity storage will be needed? Systematic modeling studies indicate that as the percentage of renewables on the grid increases, the amount of electricity storage needed to support them grows exponentially [2], but considerable disagreement remains on just how much storage is needed [2]. At the time of writing, the US consumes electricity at a rate of ≈ 500 gigawatts (GW) [3] (total US energy consumption is ≈ 3 terawatts (TW) [4]). Frew *et al.* predict that to support an 80% renewable electricity portfolio in the US, between 0.72 and 11.2 petajoules (PJ; 1 PJ = 1 × 10^15^ J or 277.8 gigawatt-hours (GWh)) of storage are needed [2, 5]. By contrast, Shaner *et al.* predict that 20 PJ of storage, about 12 hours of supply, will be needed to support 80% renewables [6]. To implement a 100% renewable electricity portfolio in the US, Frew *et al.* estimate that between 6 (without electric vehicles) and 21 (with electric vehicles) PJ of storage would be needed [2, 5, 7]. Shaner *et al.* make an even bigger prediction, that several weeks of stored supply will be needed to support 100% renewables [6]. A three-week supply of 500 GW of power amounts to 900 PJ. Projections for Europe are similar: 80% renewables need between 0.65 to 9 PJ of storage [2], while 100% requires 0.95 to 35 PJ. As economic development spreads around the world, and more and more of the global energy infrastructure is electrified (think electric vehicles) global electricity consumption will rise. Assuming that all of the 11 billion people who are projected to be alive in 2100 [8] use electricity at the rate that the average American does today (≈ 1.4 kilowatts) [9], this would correspond to a global electricity demand of ≈ 15 terawatts (TW). This may even be an underestimate, as electricity corresponds to less than 20% of US energy use per capita today [3]. Adding electrified transport into this picture could considerably increase global electricity use above 15 TW. A one-hour buffer for 15 TW would require 51 PJ (14,000 GWh) of storage, 12 hours would require 618 PJ, and three weeks would require 26 exajoules (EJ; 1 × 10^18^ J). These projected storage capacities are summarized in **Table 1**. Currently, the installed energy storage capacity in the US amounts to only ≈ 1 GWh (0.0036 PJ) [10]), while worldwide it stands at ≈ 20 GWh (0.072 PJ) [11]. How could an increase in electrical energy storage of this size be achieved?

**Table 1.** Estimated Li and Zn requirements for a representative set of energy storage scenarios.

| Scenario | Storage Requirement (Petajoules) | Amount of Li (theoretical minimum) (kilotonnes) | Amount of Li (practical) (kilotonnes) | Fraction of World Reserve | Amount of Zn (1e) (kilotonnes) | Fraction of World Reserve |
| --- | --- | --- | --- | --- | --- | --- |
| Ballpark low estimate for US or EU energy storage requirements, 80% renewables. | 1 | 19 | 47 | 0.003 | 565 | 0.002 |
| Low end estimate for 100% renewables in US, no EVs (Frew *et al.*) | 6 | 117 | 283 | 0.018 | 3,390 | 0.015 |
| Ballpark estimate for US or EU energy storage requirements, 100% renewables. | 10 | 195 | 472 | 0.030 | 5,650 | 0.025 |
| Upper end estimate for 80% renewables in US (12 hours of power) (Shaner *et al.*) | 21 | 409 | 992 | 0.064 | 11,900 | 0.052 |
| Upper end estimate for 100% renewables in US (3 weeks of power) (Shaner *et al.*) | 900 | 17,500 | 42,500 | 2.720 | 509,000 | 2.210 |
| Current world (2.5 TW), 12 hours of current supply | 108 | 2,100 | 5,100 | 0.323 | 61,000 | 0.266 |
| Current world (2.5 TW), 3 weeks of current supply | 4,540 | 88,300 | 214,200 | 13.700 | 2,560,000 | 11.200 |
| Future World, US energy consumption is standard, 11 billion people, 1 hour supply (14.3 TW) | 51 | 1,000 | 2,430 | 0.156 | 29,100 | 0.127 |
| Future World, US energy consumption is standard, 11 billion people, 12 hours supply (14.3 TW) | 618 | 12,000 | 29,200 | 1.870 | 349,000 | 1.520 |
| Future World, US energy consumption is standard, 11 billion people, 3 weeks supply | 25,900 | 505,000 | 1,230,000 | 78.500 | 14,670,000 | 63.800 |

No modern energy storage technology is perfect. Compressed air and pumped-hydro storage both have high durability [12, 13]. However, there are relatively few suitable sites for installation of either of these technologies. In addition, compressed air storage has low round trip energy storage and retrieval efficiency while the installation of pumped hydro requires a high capital investment [14]. Flow batteries scale up extremely well: their capacity is only determined by the concentration and volume of their electrolyte [14, 15]. However, current flow batteries suffer from low performance due to non-uniform pressure drops [16]. Furthermore, disposal of flow battery electrolytes poses significant environmental concerns [14]. Conventional batteries have fast response times as short as a few milliseconds [14, 17], offer an excellent combination of energy and power density for on-grid applications, and can be situated almost anywhere, making them highly scalable [18]. However, further improvements in power density in Li-batteries by decreasing the cathode thickness are limited by dendrite formation [19, 20]. The most pressing concern with all battery technologies are limited cycle and calendar lifespans. For example Li-ion batteries typically have lifespans of only 5 to 15 years or 1,000 deep charge-discharge cycles [21].

In the absence of effective recycling technologies for battery materials, the short lifespans of batteries will be significantly exacerbated by the challenges of materials availability. The total mass of electrode material, *M_electrode_* (in grams), needed to build a battery with a capacity *E_battery_* (in joules), depends on the mass of metal needed to store a unit of energy *μ_metal_* (in grams per joule),

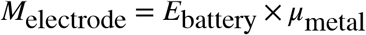

The minimum value of of *μ_metal_* can be estimated from the molecular weight of the electrolyte material (MW_metal_, in the case of Li this is 6.941), the valence state of the electrolyte (*n_e_*, in the case of Li this is 1), and the cell voltage (*V_cell_*),

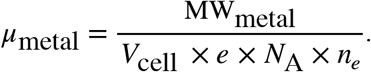

For lithium nickel manganese cobalt oxide (LiNMC; LiNiMnCoO_2_) and lithium nickel cobalt aluminum oxide (LiNCA; LiNiCoAlO_2_) cells, where *V_cell_* is 3.7 V, *μ_metal_* = 1.95 × 10^−5^ g J^−1^ (70 g kWh^−1^). In practice more than double this amount of Li is needed (≈ 170 g kWh^−1^ or 4.72 × 10^−5^ g J^−1^) [22]. Thus, in order to store 1 PJ of energy, between 19.5 and 47.2 kilotonnes of Li is required.

The total estimated masses of Li and Zn, along with the fractions of world proven reserves, needed to build the Li-ion or alkaline batteries for a wide range of projected energy storage scenarios are shown in **Table 1.** While current proven global Li and Zn reserves can easily supply the energy storage needs of Europe and the US for decades to come, should global renewable energy demand continue to rise, then global supplies of these important metals could be rapidly overwhelmed.

Many innovations will be required to allow high penetration of renewables into the global electricity supply without building a large excess of renewable capacity. New environmentally-friendly, low-cost recycling technologies for battery materials will be essential, some of which may be biological [23]. Likewise, new technologies for the synthesis of batteries at room temperature and pressure will be needed to reduce the embedded energy and carbon footprint of energy storage [24–26]. Finally, as we discuss in this article, a crucial innovation will be the development of biologically based storage technologies that use Earth-abundant elements and atmospheric CO_2_ to store renewable electricity at high efficiency, dispatchability and scalability.

## Biology Gives a First Draft Template for Storing Renewable Energy

Biology, through photosynthesis, gives a first draft template for storing solar energy at an enormous scale. Across the globe, it’s estimated that photosynthetic organisms capture solar power at an average rate of ≈ 4,000 EJ yr^−1^ (corresponding to an annually averaged rate of ≈ 130 terawatts (TW)) [27]. This energy capture rate is approximately 6.5 times greater than current world primary energy consumption of 20 TW [28]. Terrestrial photosynthetic organisms store this energy, after losses of carbon due to respiration, at a net rate of ≈ 1,200 EJ yr^−1^ (or ≈ 38 TW) largely as lignocellulosic biomass [29]. Capturing this energy requires ≈ 120 gigatonnes of carbon per year (GtC yr^−1^) (counting just the carbon atoms in fixed CO_2_) [30], while storing it requires ≈ 60 GtC yr^−1^ [31], accounting for between only 7 and 14% of global atmospheric pool of carbon [32, 33].

However, photosynthesis is far from perfect. Photosynthesis draws carbon from the atmosphere at an annually averaged rate of only 1 to 2 × 10^18^ molecules of CO_2_ m^−2^ s^−1^ [34], between 25 and 70 times less than the maximum possible uptake rate of carbon from the atmosphere of 5 to 7 × 10^19^ molecules of CO_2_ m^−2^ s^−1^ [34, 35]. As a result, the globally and annually averaged efficiency of photosynthesis ranges from between 0.25% [35] to 1% [36], with the best overall efficiencies seen in the field of between 2.4% for C_3_ plants [37], 3.4% for C_4_ plants [38] and 3% for algae grown in bubbled photobioreactors [39]. These observed efficiencies fall well below the theoretical maximum efficiencies of C_3_, C_4_, and algal photosynthesis of 4.6%, 6% [40], and 9% [39] respectively. Additionally, photosynthesis is not immediately dispatchable: it takes an entire growing season to store solar energy as plant biomass, followed by harvesting and a long series of thermochemical steps to extract energy from it.

## Components of Re-wired Carbon Fixation

### Overview

Previous analysis by us suggests that much of the inefficiency of photosynthesis arises because all of the steps of natural photosynthesis happen inside a single cell [41, 42]. Simply put, a single cell is much better at absorbing light than it is at fixing CO_2_, even when packed with the CO_2_-fixing enzyme RuBisCO. The cell absorbs far more light than it can possibly use to fix CO_2_, and dissipates the excess as heat. This leads to inefficient parallelization of the CO_2_-fixation process, and causes the efficiency of photosynthesis to drop well below its theoretical maximum [41, 42].

The rate mismatch between the light absorption and CO_2_-fixation capability in a single cell have led to attempts to rewire photosynthesis by spatially separating each of the tasks usually performed together inside a photosynthetic organism and replacing some of them with non-biological equivalents. These schemes are often called *microbial electrosynthesis,* or more recently *rewired carbon fixation.* Although originally meant to enable capture and storage of solar energy as biofuels with much higher efficiencies than photosynthesis, this separation enables the use of biology to store energy from any electrical source. A schematic of the key components of a rewired carbon fixation system is shown in **Figure 1**: sustainable energy capture (**Figure 1A**); water splitting (**Figure 1B**); electrochemical CO_2_-fixation (**Figure 1C**) and further biological reduction (**Figure 1D**) or biological CO_2_-fixation (**Figure 1E**); long-range electron transport to biological metabolism (**Figure 1F**); and energy storage molecule synthesis (**Figure 1G**). Capture of energy from sustainable energy sources (including light) (**Figure 1A**), water splitting (**Figure 1B**), and even the initial steps of CO_2_-fixation (**Figure 1C**) can now be replaced by non-biological processes, but full reduction of carbon (**Figures 1D** and **E**) and the synthesis of complex molecules (**Figure 1G**) remains exclusively the job of biology.

**Figure 1.**
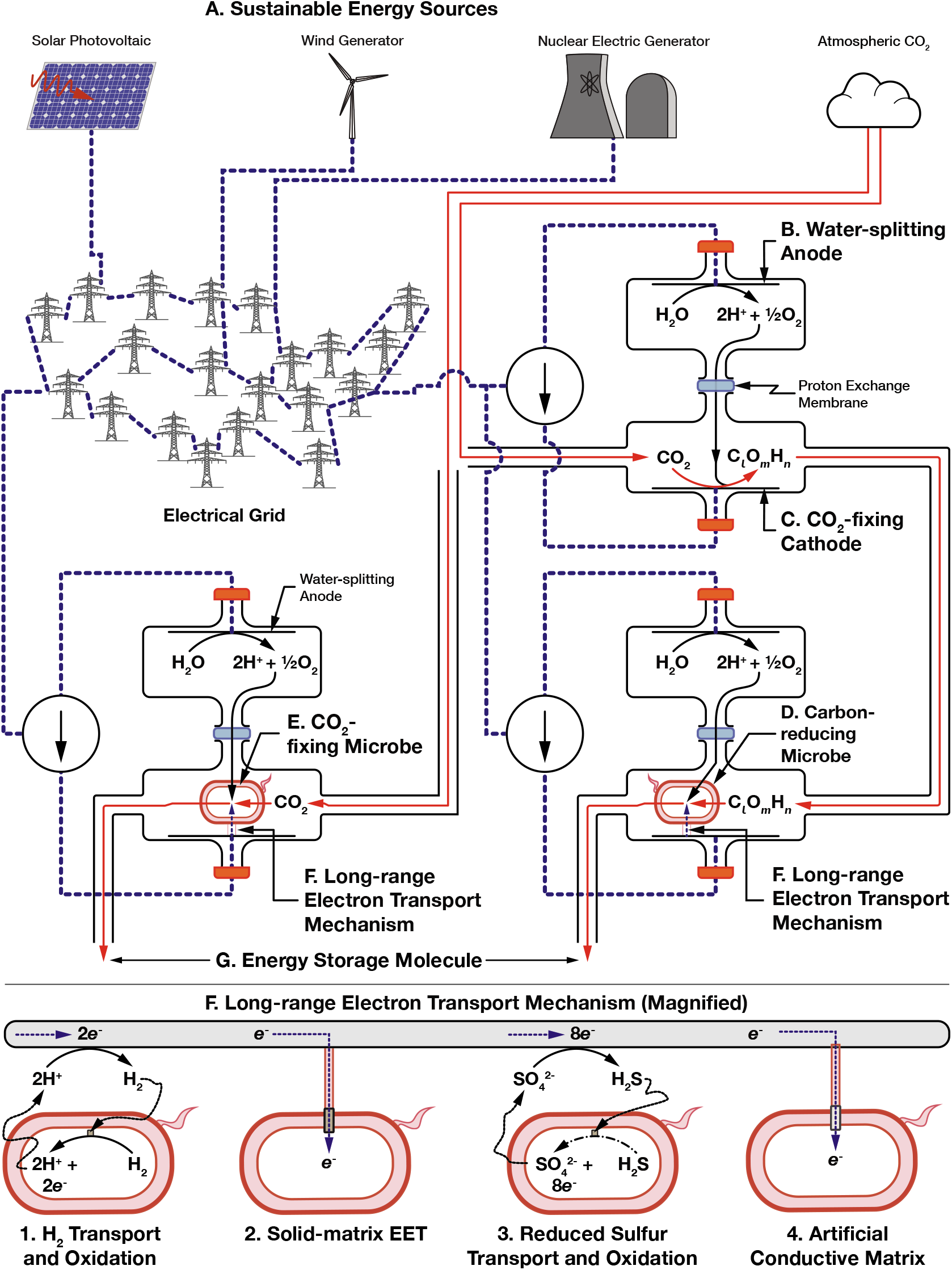
Overview of rewired carbon fixation technologies for electrical energy storage.

Several demonstrations of rewired carbon fixation have already been made, some with efficiencies exceeding that of natural photosynthesis [43–45]. However, to date, while we have previously reviewed some of constraints faced by these systems [41], no one has made a systematic review of the potential energy losses in these systems, made an upper estimate of the potential energy storage efficiency of these systems, or identified the trade-offs that the components of these systems must make. In this article, we seek to identify and catalog the parameters necessary to make this estimate, and we further identify components of the system that could be optimized by biological engineering.

#### Long-range Electron Transport and Uptake

Because rewired carbon fixation separates processes that were once performed inside a single cell, it needs mechanisms to move electrons and partially reduced carbon between components of the system that are separated by distances much longer than a single cell. Long-range electron transport and electron uptake mechanisms from non-light driven autotrophic metabolisms to move electrons from a cathode to intracellular reductants where they can be used to reduce carbon is the defining feature, and key challenge, of rewired carbon fixation. The choice of electron transfer mechanism could open up unique opportunities for the design of the system, but also set unique constraints.

The two most prominent mechanisms for long-range electron transport used in rewired carbon fixation to date are the transport of hydrogen to H_2_-oxidizing microbes [45, 46] and solid-matrix extracellular electron transfer (SmEET) enabled by conductive pili secreted by electroactive microbes [41, 47]. However, these well-known mechanisms come with a number of drawbacks including rate, safety, and poor genetic tractability. Alternative electron transport mechanisms that rely upon transport and oxidation of reduced sulfur compounds, and artificial conductive matrices could solve many of these limitations.

#### Hydrogen Transport and Oxidation

On the face of it, hydrogen has many attractive features as an electron transport mechanism for rewired carbon fixation. Its redox potential is well matched to that of NAD(P)H, the intracellular reductant used in CO_2_-fixation and many biosynthetic reactions (−0.42 V vs. the Standard Hydrogen Electrode (SHE) for 2H^+^ + 2e^−^/H_2_; and −0.32 V vs. SHE for NAD(P)^+^ + 2e^−^/NAD(P)H). It can be readily produced electrochemically with high Faradaic efficiency (> 90 % [48]) under optimized conditions, and then easily transported to a microbial culture in the gas phase; and unlike other low redox potential redox mediators like methyl viologen [49, 50] has no negative effect on microbial integrity [51].

In addition to these physicochemical advantages, H_2_ is oxidized at the cell by highly active hydrogenase enzymes that impose a very low protein load on the host cell [41]. In the H_2_-oxidizing, CO_2_-fixing microbe *Ralstonia eutropha,* H_2_ is oxidized by an inner membrane-bound hydrogenase (MBH) and a cytoplasmic soluble hydrogenase (SH). The membrane-bound hydrogenase injects electrons from H_2_-oxidation into the electron transport chain on the inner membrane, eventually reducing O_2_ and creating a proton gradient, which is used to generate ATP [52]. The soluble hydrogenase directly reduces NAD^+^ to NADH [53]. *R. eutropha* uses the ATP and NADH to fix CO_2_ through the Calvin cycle and further concatenate and reduce it to the energy storage polymer polyhydroxybutyrate (PHB) [54]. This pathway can be repurposed to produce fuels like isobutanol [43], or isopropanol [45] from electrochemically reduced H_2_.

A rewired carbon fixation system using H_2_ produced by a Co-P alloy electrode with low overpotential coupled with CO_2_-fixation and biofuel synthesis by *R. eutropha* has already achieved maximum electrical to fuel conversion efficiencies of 39%. Assuming an 18% efficient solar photovoltaic, this corresponds to a solar to fusel alcohol efficiency of 7.1% [45]. This significantly exceeds the efficiency of photosynthesis in many practical situations and almost matches the maximum theoretical efficiency of algal photosynthesis (the most efficient form of photosynthesis). However, it remains unclear how far the efficiency of this system is from its theoretical maximum, nor does a roadmap exist for achieving this efficiency, particularly through biological engineering.

The scale-up of H_2_-mediated rewired carbon fixation poses several challenges. First, in order to extract maximum energy from H_2_, O_2_ is needed as a terminal electron acceptor. This combination poses a significant explosion risk that can be mitigated by reducing the O_2_ and H_2_ concentrations in the system to below the explosive threshold (<5% H_2_), but this comes at the expense of operating rate. Secondly, many materials are highly permeable to H_2 [55]_, posing both a safety challenge and energy loss mechanism, and may even pose a risk to global climate [56]. While these safety and operational concerns can be assuaged at lab scale, it is unclear if such a system could be reliably deployed at grid-scale at a reasonable cost.

Even if these safety concerns could be circumvented, the low solubility of H_2_ in water poses a more fundamental challenge (0.0016 g/kg H_2_O or 0.8 mM for H_2_ versus 1.69 g/kg H_2_O or 38 mM for CO_2_ at 20 °C and 0.1 MPa [57]). A simple model of rewired carbon fixation mediated by H_2_ diffusion demonstrated that extremely high internal surface areas will be required for full utilization of the current produced by a 1 m^2^ solar panel [41]. This will likely require some creative engineering to maintain energy high conversion efficiency, minimize losses of H_2_, maintain acceptable safety, and prevent proton consumption due to fuel synthesis increasing solution pH to unmanageable levels [41]. While ingenious solutions to this problem do exist, such as the hollow-fiber gas reactor [58], these solutions come at the cost of high manufacturing complexity.

#### Solid-matrix Extracellular Electron Transfer and Direct Contact

At the opposite end of the spectrum of biological solutions for long-range electron transport are solid-matrix extracellular electron transfer (SmEET) mechanisms used by electroactive microbes [47]. Note, the widely accepted definition of EET does include soluble mediators like flavins [59, 60], but we do not discuss them here. These solid-matrix systems could circumvent the design challenges created by the volatility and low solubility of H_2_ in water by transferring electrons along conductive nanowires secreted by the cell, or by direct contact of the cell surface with an electrode [61].

SmEET involves three parts: long-range transport of electrons often over many cell lengths from an electrode to the cell surface; transfer of electrons from the cell surface to the electron transport chain in the inner membrane; and finally, the production of intracellular reductants that can be used in CO_2_-fixation or further reduction of partially reduced carbon. Of these three steps, the second, transfer of electrons from the outer to the inner membrane using a membrane-spanning EET complex is perhaps the best understood [62]. To our knowledge there has been only one demonstration of engineered SmEET-mediated rewired carbon fixation to date, in which a CO_2_-fixing reverse tricarboxylic acid (rTCA) cycle was enabled in the electroactive microbe *Geobacter sulfurreducens* by the addition of a gene for an ATP-dependent citrate lyase [63]. Despite this breakthrough, at the time of writing, SmEET-mediated rewired carbon fixation systems have yet to achieve the success of H_2_-mediated systems. Few, if any, organisms have been discovered that can uptake electrons, fix CO_2_, and meet the needs of the synthetic biology design-build-test loop of rapid heterotrophic growth and facile genetic modification. Furthermore, the formation of biofilms and nanowire secretion do not lend themselves to a short design-build-test loop.

The lack of a suitable naturally occurring chassis organism for SmEET-mediated rewired carbon fixation leaves the option of creating a synthetic chassis by adding SmEET, CO_2_-fixation and energy storage molecule synthesis to a highly engineerable host like *Escherichia coli, Vibrio natriegens,* or an organism with a completely synthetic genome. The *Shewanella oneidensis* Mtr complex [64] and the Calvin cycle [65] have both been separately added to *E. coli* and shown to function, although at a much lower level than in their natural hosts. Getting these systems to operate at their full potential and in concert in a synthetic host will require a much more complete understanding of the physics, chemistry and genetics of SmEET and CO_2_-fixation.

SmEET can transport electrons between sources and sinks tens to hundreds of microns from the cell surface through microbial nanowires [47, 61]. These were originally studied for electron transport out of the cell but can also move electrons into the cell. There is considerable debate about the mechanism of charge transfer in nanowires [66, 67].

A redox gradient model of conduction in electroactive biofilms has been championed by Tender, Bond and colleagues and studied most extensively in *Geobacter* biofilms [68–70], but has recently been studied in mixed community films [71]. This type of conduction relies upon long-range redox diffusion, enabled by short range electron transfer between closely spaced redox cofactors embedded throughout the conductive matrix that is composed of self-assembling protein subunits [72]. The redox gradient model of conduction was established in studies of redox polymers and hydrogels containing redox cofactors [73]. The current-voltage relationships predicted by this model have been successfully used to fit electron transport rate measurements in *Geobacter* biofilms [68, 74]. A key experimentally validated prediction of this model is the rise of film conductivity with increasing temperature [69, 70].

However, while any one of the large number of multi-heme cytochromes known to be secreted by *Geobacter sulfurreducens* could be a likely candidate for the redox cofactor used in biofilm conduction, there is no direct structural evidence of inter-heme spacing that is within the short distance (≈ 10 Å) needed for short range electron hopping needed to support electron transport at the rate seen in isolated nanowires [70]. Consequently, an alternative model for conduction in *G. sulfurreducens* biofilms has been championed by Malvankar, Tuominen, Lovely and colleagues [70, 75] that relies upon charge delocalization due to pi-stacking interactions in the *G. sulfurreducens* biofilm, similar to the conduction method in polyaniline. In contrast to the redox gradient model, this model predicts that conductivity should fall with increasing temperature [75]. However, while this predicted result has been observed by Malvankar *et al.* [75] it has not been seen by other groups [70].

A representative selection of overpotentials for SmEET-mediated systems are shown in **Table 2**. Given that the redox potential of Mtr EET complex is ≈ −0.1 V vs. SHE [76, 77], the minimum cell potential in a EET-mediated rewired carbon fixation system with a water-splitting anode is ≈ 1 V (−0.1 V - 0.82 V). The overpotentials shown in **Table 2** represent a considerable fraction of this minimum potential difference, suggesting that they could be a significant energy loss mechanism in rewired carbon fixation.

**Table 2.**
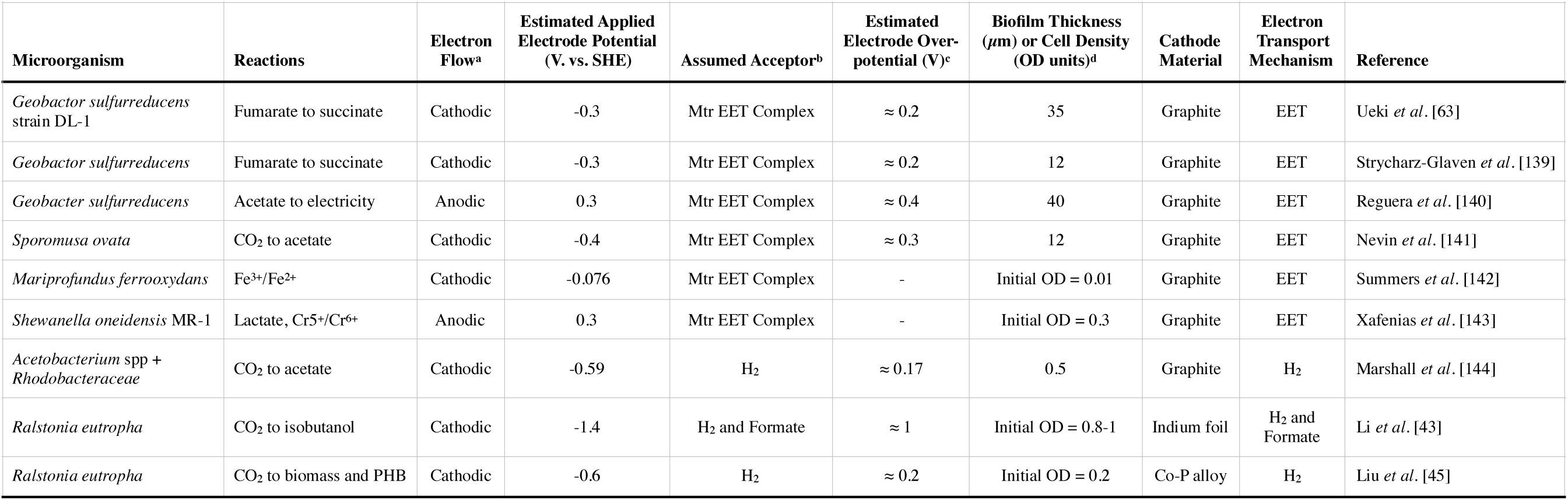
Overpotentials for a representative set of biological electron transfer systems. ^a^Cathodic electron flow refers to electron flow from cathode to microbial metabolism, whereas anodic flow indicates electron flow from metabolism towards an anode. ^b and c^The electrode overpotential is estimated by subtracting the estimated applied electrode potential from the assumed electron acceptor potential at pH 7 (Mtr EET Complex, E = −0.1 V vs SHE; H_2_, E = −0.42 V vs. SHE; Formate, E = −0.43 V vs. SHE). References [43, 45, 63, 139–144] were used to compile this table.

What is the lowest overpotential, or highest biofilm conductivity, that could be achieved? The maximum bulk *Geobacter* biofilm conductivity observed by Yates *et al.* was on the order of 5 × 10^−6^ S cm^−1^ at 30 °C (a resistivity of 2 × 10^5^ Ω cm) [69]. In contrast, Malvankar *et al.* report much higher bulk *Geobacter* biofilm conductivities of ≈ 5 × 10^−3^ S cm^−1^ (2 × 10^2^ Ω cm) [75]. The source of this discrepancy is unclear. Measurements by El Naggar *et al.* of dried isolated *S. oneidensis* nanowires indicate a resistivity on the order of only 1 Ω cm [78]. Calculations by Polizzi *et al.* suggest that such a low resistivity in a biological material could only be achieved by electron transfer with extremely closely spaced (≈ 10 Å) redox cofactors, and very low reorganization energies [72].

Gram-negative electroactive microbes have evolved an EET-complex that spans the periplasmic gap and moves electrons between the outer membrane and the electron transport chain in the inner membrane. This paradigm was first established in the electroactive microbe *S. oneidensis* MR-1, that uses the Mtr EET complex to expel electrons from metabolism onto external substrates like minerals, metal ions and even electrodes in the absence of O_2_, essentially breathing onto them [47, 79]. Similar systems containing homologous components also exist in electroactive microbes that specialize in electron uptake from metal oxidation: the phototrophic iron oxidation (Pio) complex in *Rhodopseudomonas palustris* TIE-1 [80] and *Marinobacter subterrani* [81]. While *M. subterrani* is readily genetically modifiable, it is not able to fix CO_2_. On the other hand, *R. palustris* and *S. lithotrophicus* can both fix CO_2_, but are not easily genetically modified. To our knowledge, no one has successfully coaxed *S. lithotrophicus* into forming colonies on agar, let alone grown it heterotrophically, or genetically modified it. Furthermore, Ross *et al. [82]* were able to show that the Mtr complex in *S. oneidensis* was reversible, allowing cathodically supplied electrons to catalyze the periplasmic reduction of fumarate. Measurement of the redox potentials of the *S. oneidensis* Mtr EET complex by Firer-Sherwood *et al*. [76] indicate a potential difference between the outer membrane MtrB cytochrome and the quinone pool of only about 0.0885 V, suggesting that the energy losses in this step could be much lower than in electron transport from the cathode to the cell surface.

Enabling CO_2_-fixation requires a system for generation of low-potential intracellular reductants with cathodically supplied electrons. In nature, these electrons are typically supplied to autotrophic microbes like *S. lithotrophicus* by the oxidation of Fe(II) and Fe(II)-containing minerals. This raises the issue of energetics mismatch: while the redox potential for NAD(P)^+^/NAD(P)H is −0.32 V vs. SHE [83], the redox potentials of Fe(II) and many Fe-containing minerals at circumneutral pH are several hundred millivolts higher [77]. While some Fe-oxidizing microbes like *R. palustris* [84] can use light as an additional source of energy to assist in NAD(P)^+^ reduction, others such as *M. subterrani [81]* and *S. lithotrophicus* ES-1 [80] are able to draw electrons from the oxidation of iron minerals with no external energy input.

It has long been speculated that autotrophic Fe-oxidizers use reverse electron transport to reduce NAD(P)^+^ [85]. In summary, Fe-oxidizing microbes are thought to use the EET complex to transport electrons across the periplasmic gap and into the quinone pool, at a redox potential of approximately −0.1 V vs. SHE [77]. From here the incoming stream of electrons is split into two: one stream is directed downhill in energy toward the reduction of O_2_, generating a proton gradient across the inner membrane of the cell. This proton motive force is used to generate ATP and raise the energy of the second stream of electrons to enable reduction of NAD(P)^+^. This process has been called the “uphill pathway” [77]. Recently, Rowe *et al.* [86] provided compelling evidence that cathodically supplied electrons can reduce NAD(P)^+^ in *S. oneidensis*, suggesting that this organism does indeed contain such a pathway.

Should the existence of the uphill pathway in *S. oneidensis* be confirmed, two immediate questions are raised: what are the components of this pathway, and how is electron flow between the uphill and downhill branches of the pathway regulated? Furthermore, if the components of this pathway could be isolated and used in rewired carbon fixation, what costs does this system impose on overall system efficiency?

#### Sulfur Transport and Oxidation

The limitations of hydrogen transport and SmEET have inspired searches for alternative mechanisms of long-range electron transport. Several choices have been proposed that can be renewably sourced including ammonia (NH_3_), phosphite (HPO_3_-), and reduced sulfur compounds (H_2_S, S_2_O_3_^2-^, S_4_O_6_^2-^) [87]. While ammonia has high solubility in water, its metabolic oxidation product NO_2_-has high microbial toxicity [87]. Phosphite and its oxidation product phosphate (PO_4_^3-^) have low toxicity, and both are highly soluble in water. However, the use of phosphite as a redox mediator comes with a potentially large energy loss. The phosphite/phosphate couple has a redox potential of −0.65 V vs. SHE. However, phosphite directly donates electrons to NAD(P)^+^ through phosphite dehydrogenase, leading to an overpotential loss of over 300 mV [88].

Sulfur can be found in nature in a wide range of oxidation states, from −2 and up to 6, allowing it to carry up to 8 electrons per atom. Each of these oxidation states, except for the most oxidized, can be used as an electron donor for chemoautotrophic microbial growth. The most common sulfur compounds used as electron donors are hydrogen sulfide (H_2_S), elemental sulfur (S^0^), tetrathionate (S_4_O_6_^2-^), and thiosulfate (S_2_O_3_^2-^) [89]. Each of these compounds can be microbially oxidized to sulfate (SO_4_^2-^) [89]. Reduced sulfur compounds (with the exception of S^0^) are far more soluble in water than hydrogen (2.5g/kg H_2_O or 110 mM for H_2_S, 1.4 M for Na_2_S_2_O_3_, and 113 mM for Na_2_S_4_O_6_, versus 0.8 mM for H_2_ at 20 °C) [90]. Given that diffusional transfer rate increases with mediator concentration, this has the potential to dramatically increase rates of energy and charge transfer to metabolism, and reduce the internal complexity of the electrosynthesis reactor [41]. As reduced sulfur compounds transfer electrons by diffusion, rather than relying upon a solid matrix, they are suitable for the rapid design-build-test cycle used in synthetic biology. On top of this, hydrogen sulfide, thiosulfate and tetrathionate are far less volatile and flammable than hydrogen, significantly reducing operational safety concerns [91].

It is now possible to electrochemically recycle sulfate, enabling a continuous transfer of electrons to microbial metabolism from a cathode. Bilal and Tributsch demonstrated reduction of sulfate to sulfide on graphite electrode at an applied potential of 1.5 V vs. SHE, with a bias of 1 V, at temperatures close to 120 °C [92]. Sulfate can also be directly reduced to tetrathionate at an applied potential of ≈ 1.7 V vs. SHE on a vitreous carbon electrode [93, 94]. While electrochemically reducing sulfate directly to thiosulfate is difficult at lab scale due to the high Gibbs free energy of this reaction (*ΔG* ≈ 700 kJ mol^−1^) [95], it is conceivable that this reduction could be catalyzed by multiple reduction steps [96, 97].

Sulfur-oxidizing microbes are often found in the mixing zone between oxygenated seawater and reduced hydrothermal fluids in the vicinity of deep-sea hydrothermal vents. Free-living species including *Thiomicrospira* and *Beggiatoa* are found above the seafloor [98], while species like *Sulfurimonas* are found below it [99]. Amazingly, sulfur-oxidizing microbes are often found inside invertebrates living near hydrothermal vents, providing them with sugar produced directly from carbon dioxide dissolved in the seawater [99–101].

Two pathways for sulfur oxidation are known that enable microorganisms to oxidize reduced sulfur compounds including hydrogen sulfide (**Figure 2**), tetrathionate (**Figure 3**), and thiosulfate (**Figure 4**) to sulfate and use the extracted energy and charge to power chemoautotrophic metabolism. In the Sox (sulfur oxidation) system (**Figures 2A, 3A** and **4A**), first established in studies of *Paracoccus pantotrophus* and *Sulfurimonas denitrificans,* reduced sulfur compounds are immobilized on the SoxY protein and repeatedly oxidized by the SoxCD protein, before final oxidation to sulfate by SoxB [102, 103].

**Figure 2.**
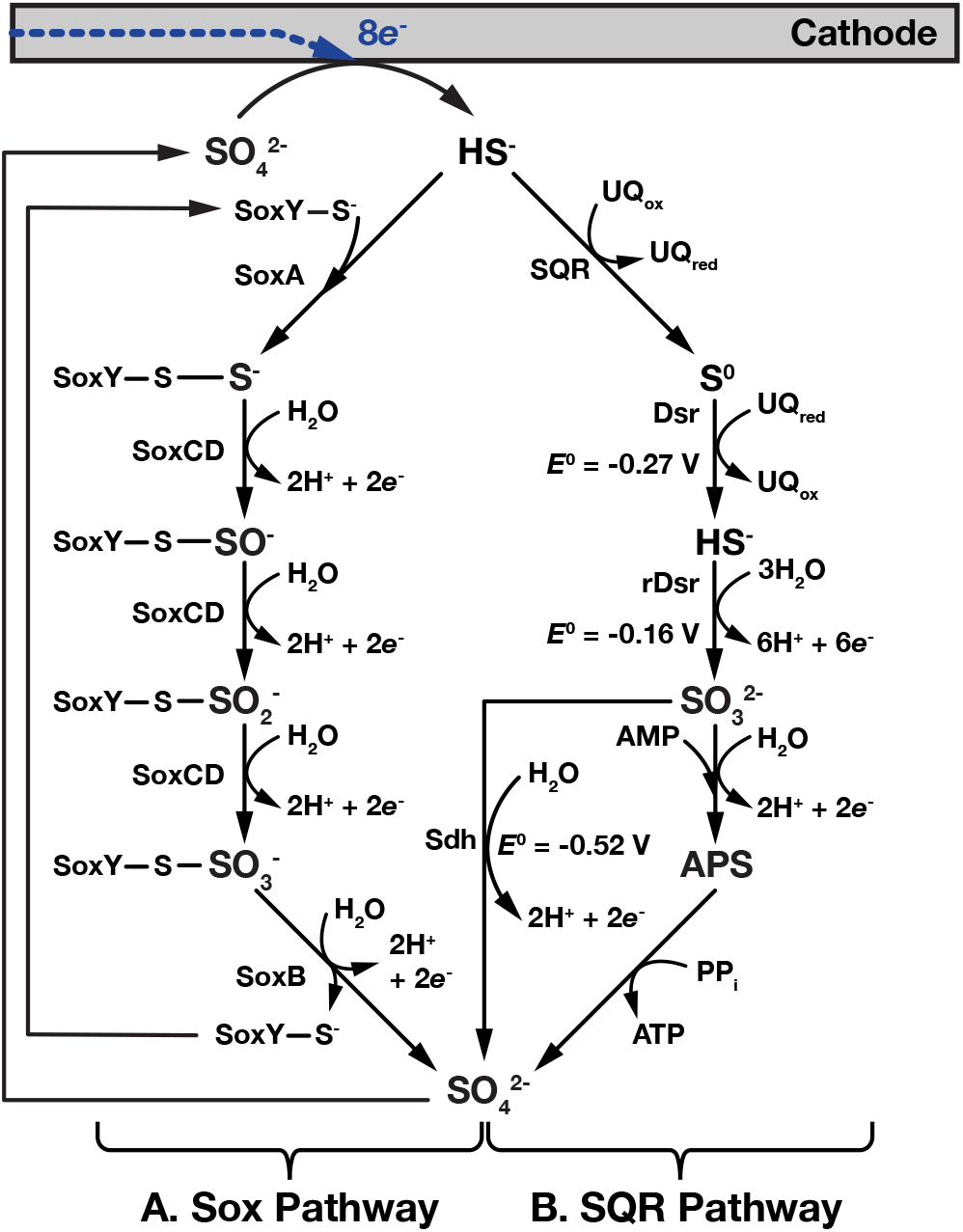
Enzymatic pathways for oxidation of electrochemically reduced hydrogen sulfide. In the Sox (Sulfide oxidation) pathway (**Figure 2A**), located in the periplasm of the microorganism, sulfide is bound to the SoxY enzyme through a cysteine-sulfur atom (SoxY-S^-^) and is sequentially oxidized to sulfate. SoxCD is believed to catalyze the oxidation through to sulfite (SO_3_-), with the final oxidation to sulfate (SO_4_^2-^) catalyzed by SoxB. The sulfide quinone oxidoreductase (SQR) pathway (**Figure 2B**), includes the formation of the free intermediates elemental sulfur (S^0^), sulfite (SO_3_^2-^) and APS (adenosine-5′-phosphosulfate). In this pathway, hydrogen sulfide is first oxidized to sulfur in a 2-electron reaction by a sulfide:quinone reductase (SQR). In *Beggiatoa* this sulfur precipitates and is stored in intracellular granules. When the supply of sulfide has been depleted, elemental sulfur can be converted back to soluble sulfide and sent to the cytoplasm by the Dissimilatory sulfite reductase (Dsr), a membrane spanning siroheme. Sulfide is further oxidized to sulfite by reverse Dsr (rDsr), then to sulfate by either APS reductase and ATP sulfurylase, or Adenosine 5’-monophosphate (AMP)-independent sulfite dehydrogenase (Sdh). This cycle is completed when sulfate is electrochemically reduced back to sulfide at the cathode. This figure was compiled with information from references [103, 104, 137, 138].

**Figure 3.**
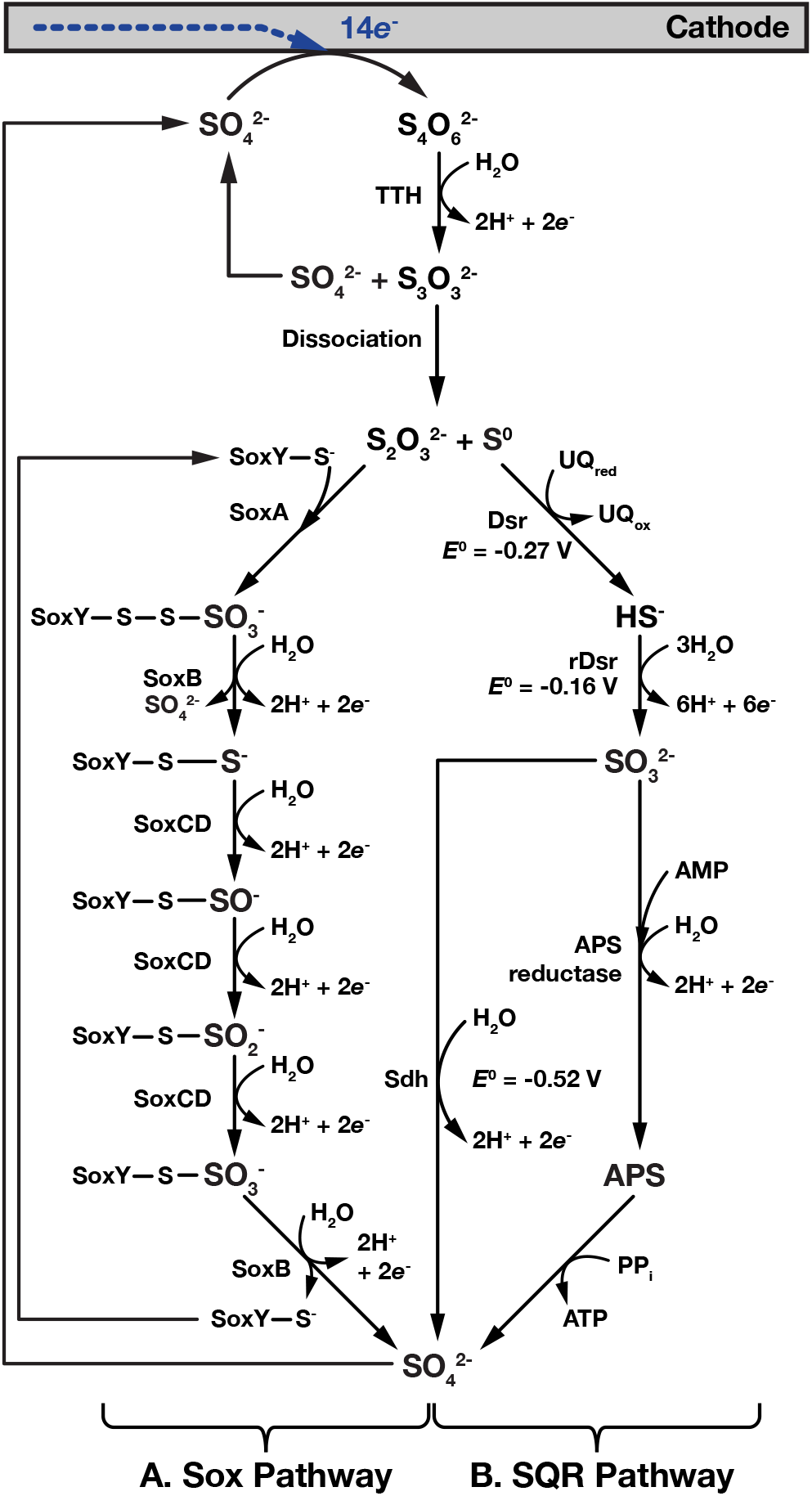
Enzymatic pathways for oxidation of electrochemically reduced tetrathionate. Tetrathionate (S_4_O_6_^2-^) is oxidized by a membrane-bound Tetrathionate hydrolase (TTH) to sulfate and thioperoxymonosulfate (S_3_O_3_^2-^) which spontaneously dissociates into sulfur (S^0^) and thiosulfate (S_2_O_3_^2-^). Elemental sulfur is converted to sulfide by the Dissimilatory sulfite reductase (Dsr), then following the pathway shown is **Figure 2B**, sulfide is oxidized to sulfate. Thiosulfate is oxidized via the Sox pathway, similar to that shown in **Figure 2A**. However, an additional oxidation step, catalyzed by SoxB at the beginning of the pathway, releases an additional sulfate molecule, that can also be recycled back to tetrathionate via cathode reduction. This cycle is completed when sulfate is electrochemically reduced back to tetrathionate at the cathode.

**Figure 4.**
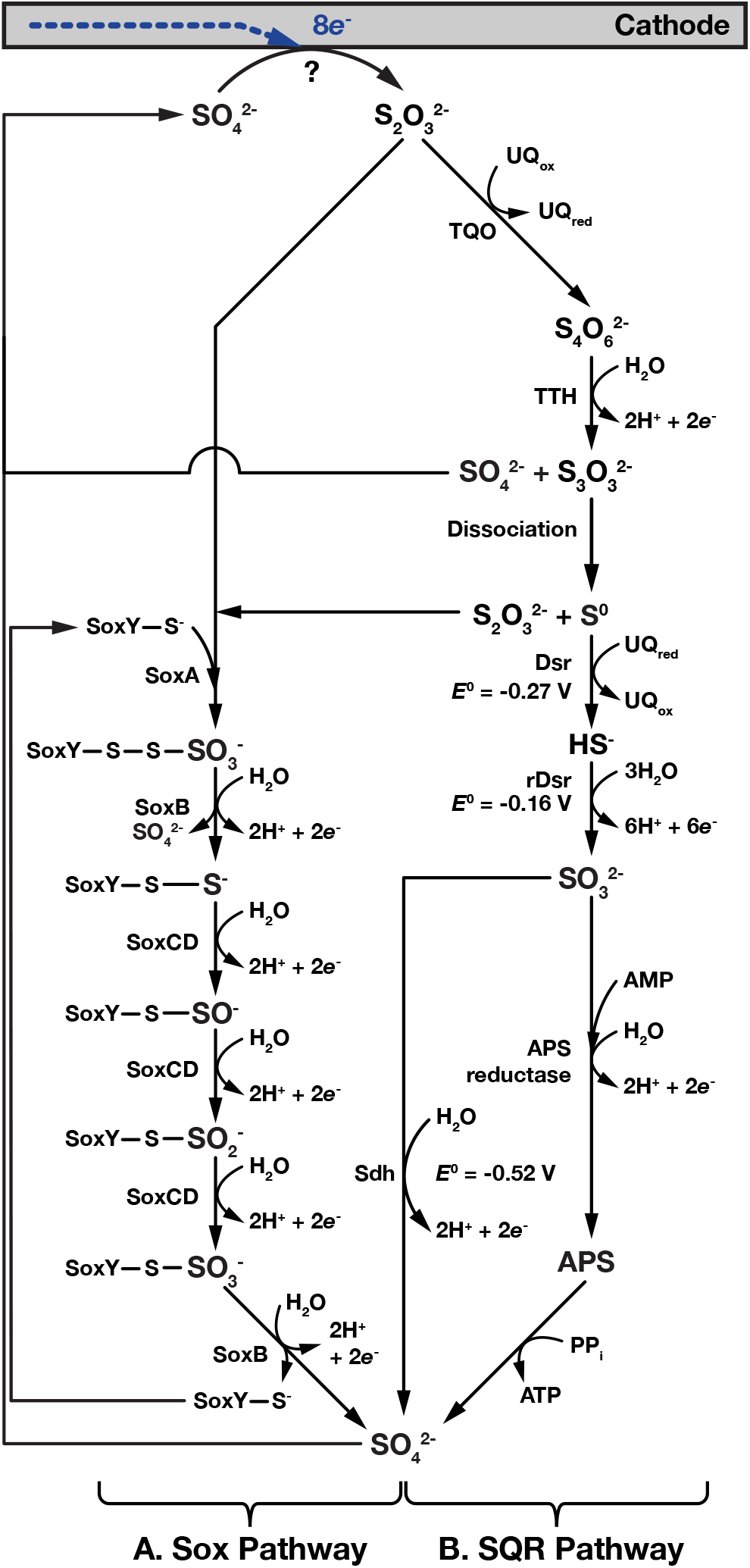
Enzymatic pathways for oxidation of electrochemically reduced thiosulfate. Although challenging, it may be possible to electrochemicallyreduce sulfate to thiosulfate (we have placed a ? at the sulfate to thiosulfate reaction to indicate this difficulty). Thiosulfate can be directly oxidized to sulfate through the Sox system (**Figure 4A**). Also, a membrane-bound, the thiosulfate:quinone oxidoreductase (TQO) can oxidize thiosulfate to tetrathionate in a 2-electron reaction (**Figure 4B**), followed by oxidation to sulfate through the tetrathionate oxidation pathways shown in **Figure 3**. This cycle is completed when sulfate is electrochemically reduced back to thiosulfate at the cathode.

The oxidation of reduced sulfur compounds can also occur through a series of non-immobilized intermediates through the full sulfide:quinone oxidoreductase (SQR) pathway (**Figure 2B**) or parts of it (**Figures 3B** and **4B**). When beginning with H_2_S, microorganisms such as *Thiobacillus denitreficans* and *Beggiatoa* first use the sulfide:quinone oxidoreductase to oxidize H_2_S to insoluble elemental sulfur (S^0^) that accumulates in the cell’s periplasm [104]. When the supply of sulfide has been depleted the stored sulfur is first reduced to HS-by the periplasmic Dissimilatory sulfite reductase (Dsr), followed by a 6-electron oxidation to sulfite at a redox potential of −0.16 V vs. SHE by the reverse Dissimilatory sulfite reductase (rDsr) [89] [95]. Finally, sulfite is oxidized to sulfate with the release of two electrons (**Figure 2B**).

The first step of the SQR pathway can be bypassed to enable oxidation of tetrathionate (S_4_O_6_^2-^), and thiosulfate (S_2_O_3_^2-^) (**Figures 3B** and **4B**). Tetrathionate is first oxidized by Tetrathionate hydrolase (TTH) to sulfate and thioperoxymonosulfate (S_3_O_3_^2-^ Thioperoxymonosulfate then dissociates to thiosulfate and elemental sulfur that are oxidized by the Sox pathway and post SQR steps of the SQR pathway respectively (**Figures 3A** and **B**).

Thiosulfate is first oxidized by thiosulfate:quinone oxidoreductase (TQO) to tetrathionate which is then then oxidized by TTH to produce sulfate and thioperoxymonosulfate. As before, thioperoxymonosulfate then dissociates to thiosulfate and elemental sulfur that are oxidized by the Sox pathway and post SQR steps of the SQR pathway respectively (**Figures 4A** and **B**).

In all sulfur oxidation pathways the starting substrates are oxidized to sulfite before final oxidation to sulfate. While the reduction potential of sulfite/sulfate is very low (E = −515 mV vs. SHE) [83], at the time of writing, we are unaware of any reports of an enzyme that catalyzes the transfer of electrons from sulfite to NAD(P)^+ [87]^. Therefore, the microbial utilization of reduced sulfur species is thought to involve reverse electron flow (also known as the uphill pathway). Were sulfur oxidation to be used in rewired carbon fixation, the effect of use of reverse electron flow on the efficiency of the system is unknown. However, use of reverse electron flow does possibly avoid the overpotential losses seen in phosphite oxidation.

In addition to the desirable physicochemical properties of reduced sulfur compounds, this mode of long-range electron transport also comes with biological advantages. Each of the sulfur oxidation pathways presented here are composed of a large number of genes, many of which are known, making reconstitution in heterologous hosts like *E. coli* or *V. natriegens* challenging but almost certainly possible. Furthermore, the large number of organisms that use sulfur-oxidation exist in a wide range of environments with differing pH and temperature [105]. This gives us a large selection from which to find an easily genetically tractable organism which can be characterized to find the full set of genes needed for sulfur-oxidation and possibly one that meets the needs of the synthetic biology design-test-build loop, and a fully operational rewired carbon fixation system.

#### Artificial Conductive Matrices

The limitations of naturally occurring electroactive biofilms both during the prototyping phase of synthetic biology and later during application could be addressed by building artificial conductive matrices tailored for rewired carbon fixation.

Recent works demonstrate that non-biologically synthesized conductive matrices can enhance power output in microbial fuel cells. Yu *et al.* [106] developed an artificial conductive matrix composed of graphite particles wrapped in conductive polymer chains of polypyrrole. A microbial fuel cell using *S. oneidensis* embedded in this artificial matrix produced 11 times more power than a comparable cell using a natural *S. oneidensis* biofilm. Estevez-Canales *et al.* [107] developed an artificial conductive matrix for *G. sulfurreducens* composed of carbon felt fibers embedded in silica gel. The silica-carbon composite allowed rapid encapsulation of *G. sulfurreducens,* which could allow for rapid prototyping of engineered electroactive microbes in the lab. However, neither of these approaches are amenable to self-assembly and more importantly self-repair, that would allow a rewired carbon fixation system to maintain itself over long periods of time.

Recent advances in the computational design of protein molecules that self-assemble into extended structures open the possibility of creating a synthetic biological conductive matrix. Gonen *et al.* [108] designed protein homo-oligomers that could self-assemble into 2D protein arrays with a maximum thickness of 3 to 8 nm, with a maximum length of 1 μm [108]. Meanwhile, Shen *et al.* designed protein monomers that could self-assemble into filaments that were multiple μm in length [109].

A synthetic biological conductive matrix could be engineered to test the competing theories of conduction in natural biofilms and improve upon the conductivity of naturally occurring conductive biofilms in order to minimize energetic losses in rewired carbon fixation. One design class could test the redox gradient model of conduction seen in *Geobacter* biofilms. This class of conductive matrix could be engineered with embedded closely-spaced (<10 Å) metal ligands [110] that act as redox cofactors to enable long distance redox diffusion. An alternative class of design could test the organic metal model of conduction. This class of design could be engineered to contain aligned pi-stacking interactions to enable charge delocalization. If, as Polizzi *et al.* speculate [72], the conductivity of individual nanowires is already highly optimized (isolated *S. oneidensis* nanowires already have a conductivity as high as 1 S cm^−1^ [78]), considerable improvements in bulk conductivity could still be made (*G. sulfurreducens* films have a conductivity of between (5 × 10^−3^ S cm^−1^ [69] and 5 × 10^−6^ S cm^−1^ [75]) by increasing the packing density of nanowires in a conductive matrix. Further in the future, it may be possible to design a complementary synthetic conductive matrix and synthetic EET complex with redox potentials well matched to that of NAD(P)H, permitting direct reduction without the need of an uphill pathway.

### In the Cell Carbon Fixation

Room temperature and pressure, free-air carbon fixation to carbohydrates and hydrocarbons driven by light-activated water-splitting or from inorganic electron donors like Fe(II), H_2_, and reduced sulfur compounds is one of the most attractive features of biology. While *R. eutropha* is a highly attractive chassis organism for H_2_-mediated rewired carbon fixation as it contains both H_2_-oxidation and CO_2_-fixation ability, the lack of CO_2_-fixing ability in many of the most engineerable organisms for rewired carbon fixation, like *E. coli, V. natriegens,* and completely synthetic organisms, raises the need to add it. Given a large choice of naturally evolved CO_2_-fixation pathways and a growing number of proposed and even implemented synthetic alternatives (**Table 3**), this raises the choice of which one to add.

**Table 3.**
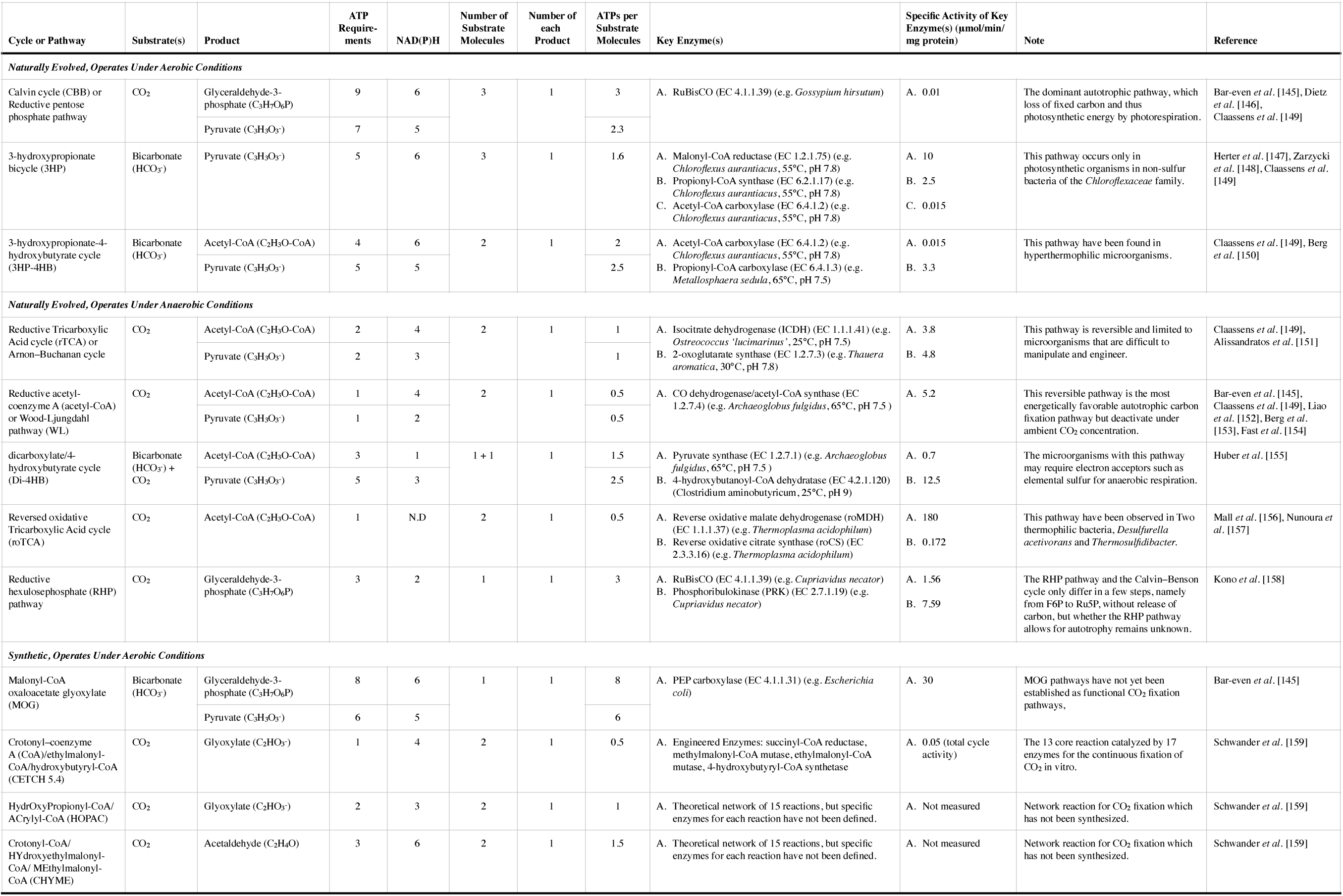
Natural and synthetic biological carbon fixation cycles and pathways. References [145–159] were used to compile this table.

In an integrated system like natural photosynthesis, where CO_2_-fixation and light capture are performed in the same cell, the photon supply can exceed the maximum possible photon utilization rate [41, 111]. This means that given the choice between thermodynamic efficiency and rate of CO_2_-fixation, evolution will likely trade some efficiency for fixation rate, as there is often an ample supply of photons.

On the other hand, in a separated system like rewired carbon fixation the overall CO_2_-fixation rate can be increased by connecting more cells. This means that the more efficient the long-range electron transport system is, the more the choice of CO_2_-fixation method can shift from one that is fast towards one that is thermodynamically efficient.

The most natural first choice of carbon fixation mechanism to engineer into a rewired carbon fixation chassis is the Calvin-Benson-Bassham cycle (CBB; or Calvin cycle) (**Table 3**). The Calvin cycle is the predominant mode of carbon fixation used in nature and is by far the best characterized. Several attempts of increasing complexity and success have been made at adding part or all of the Calvin cycle to *E. coli* to transform it into an autotroph. Most recently, Antonovsky *et al*. [65] demonstrated the synthesis of sugars from fixed carbon with the Calvin Cycle in *E. coli,* but were unable to accumulate biomass. However, despite these advantages, the Calvin cycle has high ATP and reductant (Ferredoxin and NAD(P)H) requirements per substrate molecule, and slow pathway kinetics (**Table 3**) due mainly to the poor catalytic performance of its carboxylase: RuBisCO. Aside from its slow CO_2_ fixation rate, RuBisCO also has an undesirable side-reaction with O_2_, producing one molecule of glycolate-2-phosphate (G2P) and one molecule of 3-phosphoglycerate, instead of two molecules of 3-phosphoglycerate. Recycling G2P by photorespiration releases CO_2_ and requires ATP and NADPH. Under current atmospheric CO_2_ concentrations and at 25 °C, photorespiration raises the minimum quantum requirement of C_3_ photosynthesis from 8 to 13 photons per CO_2_ assimilated [112]. It is estimated that up to 30% of the photosynthetic output is lost through photorespiration [113]. Some organisms that employ the Calvin Cycle minimize energetic losses due to photorespiration by using CO_2_-concentrating mechanisms such as bundle sheath cells in C_4_ plants and carboxysomes in cyanobacteria [114].

Given these limitations, other carbon fixation cycles found in nature could be attractive (**Table 3**). It is conceivable, given recent advances in compartmentalization in synthetic biology [115, 116] that highly efficient pathways like the Wood-Ljungdahl pathway that require high CO_2_ concentrations could be implemented under atmospheric CO_2_ concentrations in rewired carbon fixation organisms using synthetic carbon concentrating compartments or heterologously expressed carboxysomes [117].

Finally, the limitations of naturally occurring carbon fixation cycles and pathways have led to efforts to design artificial carbon fixation mechanisms with higher kinetic rates and efficiencies than natural mechanisms through new combinations of naturally occurring and synthetic enzymes. A representative set of promising synthetic cycles is shown in **Table 3**.

Implementing CO_2_-fixation in a non-native host remains a grand challenge in synthetic biology, but considerable progress has been made in the last decade. Future breakthroughs in this area could be made with better tools for the evolution of autotrophic, CO_2_-fixing organisms, and better systems biology tools to understand the genomes of heteroautotrophs like *R. eutropha* and *Chlamydomonas reinhardtii [118].*

### Out of the Cell Carbon Fixation, Transportation and Uptake

#### Overview

Recent advances in electrochemistry have enabled the reduction of CO_2_ to C_1_, C_2_ and C_3_ compounds (**Figure 1C**). A representative set of electrochemical CO_2_ reductions are shown in **Table 4**. Electrocatalysts can reduce CO_2_ to C_1_ compounds like formate and carbon monoxide with very Faradaic efficiencies and at very high rates [48]. However, the electrochemical production of higher chain length products is much more challenging [119]. Paris *et al.* [120] recently transformed CO_2_ into propanol (C_3_H_8_O) with a thin film Ni3Al electrode poised at −1.18 V vs. SHE but with a Faradaic efficiency of only 1.9 ± 0.3% (**Table 4**). The high efficiencies and rates of electrochemical conversion of CO_2_ to short chain length products, but the difficulty in conversion to higher molecular weight products, allows a process that was once exclusively performed by biology to be replaced, leaving biology to do what it does exclusively best, the highly efficient synthesis of complex carbon-containing molecules at room temperature and pressure (**Figures 1D** and **1G**).

**Table 4.**
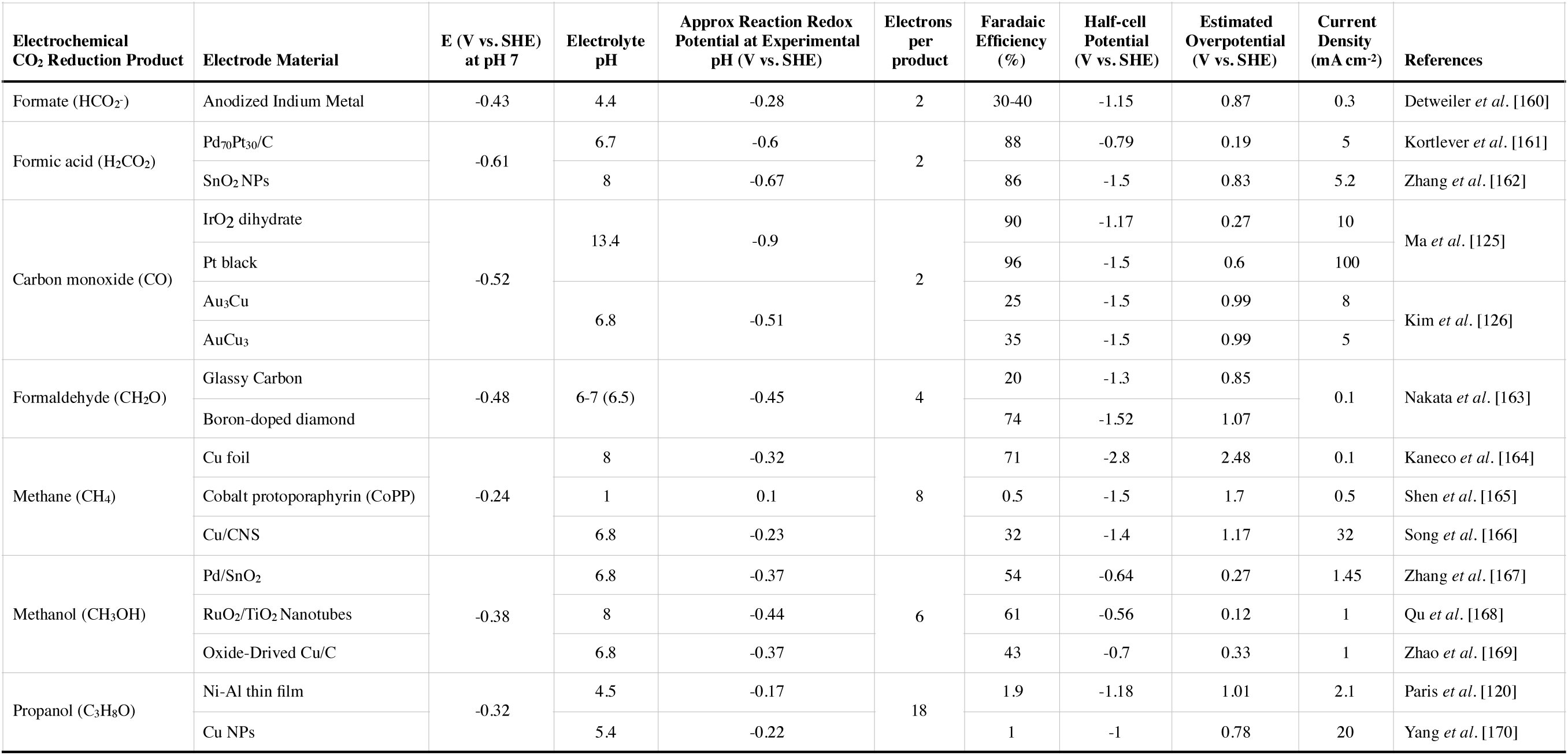
Representative set of electrochemical CO_2_ fixation schemes. This table was compiled from information in references [120, 125, 126, 160–170].

Long-range electron transport and electrochemical CO_2_ reduction are highly complementary. While microbial metabolism can concatenate and further reduce short chain carbon-containing molecules, this comes with two complications. First, in order to further reduce short chain hydrocarbons (the primary fixation molecule), the release of CO_2_ is typically required to enable the concentration of the limited number of input electrons. For example, in order to make a single PHB monomer (C_4_H_8_O_3_), a microbe would need 42 electrons (*n_e, s_*; where *s* stands for storage molecule) and 4 carbon atoms (*n_c, s_*). To source these from formate (HCO_2_-) which carries 1 carbon atom (*n_c, p_*; where *p* stands for primary fixation molecule) and 2 electrons per molecule (*n_e, p_*; where *p* stands for primary fixation molecule), the microbe would need to expend 21 formate molecules, and then re-emit 17 CO_2_ molecules, a loss of ≈ 80% of the initially fixed carbon back into the atmosphere. In principle, a carbon-reducing electroactive microbe (**Figure 1D**) could simply source the extra electrons (*n_e,add_)* to supplement the electrons carried by the primary fixation molecule from long-range electron transport to perform an unbalanced reduction,

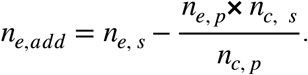

For instance, with 4 formate molecules, an electroactive microbe could in principle make one PHB monomer by absorbing an additional 34 electrons, with no re-release of carbon back into the atmosphere.

Nature provides a toolkit of enzymes and pathways for processing electrochemically reduced carbon molecules that can potentially work in concert with electron uptake. A summary of a representative set of these pathways is shown in **Table 5**.

**Table 5.**
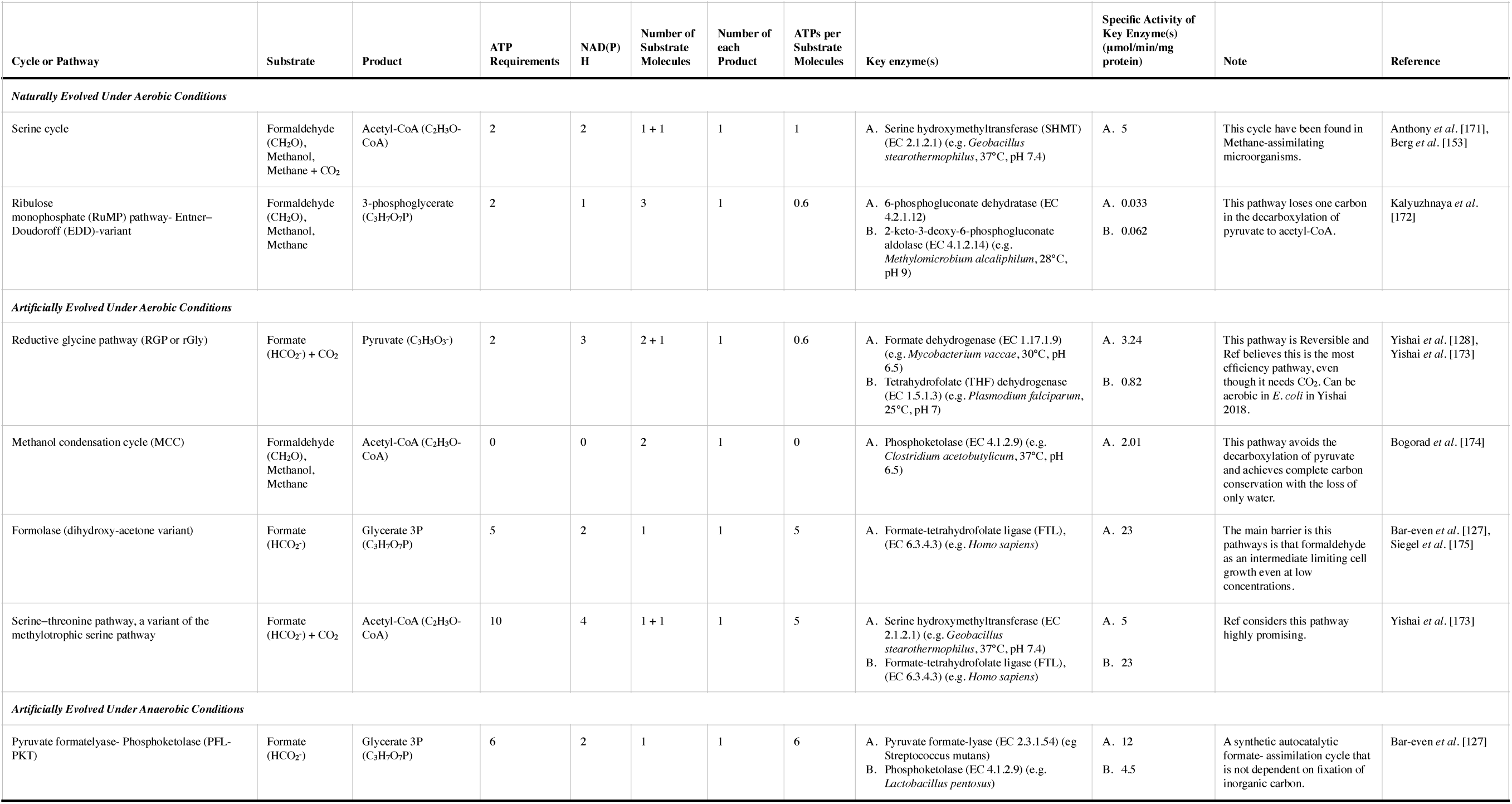
Representative set of pathways for processing partially reduced carbon. References [127, 128, 153, 171–175] were used to build this table.

#### Carbon Monoxide

Carbon dioxide can be electrochemically reduced to carbon monoxide (CO) at a redox potential of −0.52 V vs. SHE at pH 7.0 with extremely high current densities and Faradaic efficiencies as high as 96% (**Table 4**). Carbon monoxide dehydrogenase (CODH) catalyzes the reversible oxidation of CO to CO_2_, enabling growth on CO and possibly synthesis of energy storage molecules. Two classes of CODH exist: the first class is found in aerobic microbes such as *Oligotropha carboxidovoran* [121]: while the second is found in anaerobic microbes including *Moorella thermoacetica* [122], *Rhodospirillum rubrum [123],* and *Carboxydothermus hydrogenoformans [124].*

Despite these attractions, carbon monoxide has a low solubility in water (0.028 g/kg H_2_O or 1 mM), comparable to that of H_2_ (0.8 mM), approximately 100 to 1000 times lower than reduced sulfur compounds, and ≈ 45 times less soluble than CO_2_ (45 mM) [57, 90]. In addition, electrochemical reduction of CO with high Faradaic efficiency requires rare metal catalysts (Pt and Ir [125]) or nanostructured catalysts [126]. Finally, CO is flammable and highly toxic to both humans and microbes [51, 87]. Taken together, these constraints make CO far less attractive than reduced sulfur compounds, SmEET or even H_2_.

#### Formate and Formic Acid

Carbon dioxide can be electrochemically reduced to formate (HCO_2_-) at high Faradaic efficiency under circumneutral conditions (**Table 4**). In comparison to other C_1_ compounds such as methane (−0.24 V vs. SHE at pH 7.0) and methanol (−0.38 V vs. SHE at pH 7.0) [119] the low redox potential of formate (−0.42 V vs. SHE at pH 7.0) allows the direct reduction of NAD(P)^+^.

Furthermore, formate is much more soluble in water (Sodium formate has a maximum solubility of 972 g/kg H_2_O at 20 °C or 14.3 M) than methane (0.025 g/kg H_2_O at 20 °C or 1.4 mM) [90]. Li *et al.* demonstrated the production of isobutanol from electrochemically reduced formate using a synthetic pathway in *R. eutropha* [43]. However, this pathway relies upon the conversion of formate back to CO_2_ in the cell, forcing this system to be reliant upon the Calvin Cycle and all of its limitations [43]. In addition, there are several naturally occurring formate assimilation pathways that do not rely upon RuBisCO, however, at the time of writing there are no known formate assimilation pathways that do not rely upon the enzymatic incorporation of CO_2_ [127]. This means that most carbon incorporated into metabolism has to come through enzymatic routes and does not fully leverage the advantages of electrochemical reduction of CO_2_ to formate. However, recent advances in computational design of synthetic metabolic pathways have yielded several designs that do not rely upon any enzymatic fixation of CO_2_ [127, 128]. The most promising are shown in **Table 5**.

The main barrier to the use of formate as a microbial feed-stock is its toxicity to many of the bacteria that can oxidize it. Formate inhibits growth at concentrations of tens of mM by inhibiting cytochrome *c* oxidation [129] and acidifying the cytoplasm, dissipating the proton motive force [130, 131]. A major opportunity in biological engineering is to develop a rewired carbon fixation chassis organism with a higher tolerance to formate, allowing it to take full advantage of the high solubility of both reduced sulfur compounds and formate.

### Metabolism and Energy Storage

At the time of writing, rewired carbon fixation projects have focused on the production and secretion of liquid fuels for transportation. Biology offers a large selection of enzymes and complete metabolic pathways that can produce a large set of fuel molecules at room temperature and pressure including isobutanol [132], octanol [133], branched-chain alcohols [134], medium-chain fatty acids [135], and alkanes [136]. The production of transportation fuels faces several constraints, some of which are set by the physical demands of the application like high energy density and low volatility as in aviation, but also by the need for compatibility with legacy use (think engines and jet turbines), distribution and regulatory infrastructures.

However, far less attention has been paid to the synthesis of carbon-containing molecules that are tailored for the storage and retrieval of electrical energy. As this application is completely new, the constraints of this application can be largely physical in nature: energy density; non-bio-toxicity; non-volatility; and environmental safety. A promising candidate for this role are bio-plastics. Several wild-type CO_2_ fixing organisms are able to accumulate large quantities of the bioplastic polyhydroxybutyrate (PHB) within the cell. *R. eutropha* is a prolific PHB producer, can accumulate 15g-PHB per liter of culture per hour when grown on CO_2_, H_2_ and O_2_, and PHB can account for up to 87% of cell weight. Energy could be retrieved from PHB either by metabolic oxidation, and subsequent release of energy directly back to electricity through EET. Alternatively, the accumulated biomass could be gasified, and directedly converted back to electricity in a fuel cell.

## Conclusions

Biology, and particularly rewired carbon fixation, could hold the answer to the large-scale storage of renewable energy. Several key challenges must be addressed: finding a mechanism for long-range electron transport that is efficient, supports high transfer rates, safe, and can be rapidly engineered; a mechanism of carbon fixation that can be expressed in a heterologous host, and is thermodynamically highly efficient, if not also fast; and finally, an energy storage system that is safe, convenient, and enables rapid dispatchibility. These innovations will require breakthroughs in systems biology of non-model exotic microorganisms, mining the genomes of exotic organisms, evolution tools for autotrophic metabolisms and in the development of synthetic enzymes and self-assembling and self-repairing biological nanostructures.

## List of Abbreviations

EET: - Extracellular Electron Transfer
SmEET: - Solid-matrix Extracellular Electron Transfer
SHE: - Standard Hydrogen Electrode
TW: - Terawatt (1 × 10^12^ Watts)
GW: - Gigawatt (1 × 10^9^ Watts)
PJ: - Petajoule (1 × 10^15^ Joules)
EJ: - Exajoule (1 × 10^18^ Joules)
GWh: - Gigawatt-hour (3.6 petajoules)
kWh: - kilowatt-hour (3.6 megajoules)
GtC: - gigatonnes of carbon (counting just the mass of carbon atoms in a carbon compound like CO_2_)
Sox: - Sulfur oxidation system
SQR: - Sulfide Quinone Oxidoreductase
APS: - Adenosine 5′-Phosphosulfate
AMP: - Adenosine 5′-Monophosphate
TTH: - Tetrathionate Hydrolase
TQO: – Thiosulfate Quinone Oxidoreductase
Dsr: - Dissimilatory sulfite reductase
rDsr: - reverse Dissimilatory sulfite reductase
UQ_ox_: – Ubiquinone oxidation
UQ_red_: – Ubiquinone reduction
*n*_e, s_: - Number of electrons for storage molecule
*n*_c, s_: - Number of carbons for storage molecule
*n*_e, s_: - Number of electrons for primary fixation molecule
*n*_c, s_: - Number of carbons for primary fixation molecule
*n*_e, add_: - Number of needed extra electrons.

## Declarations

### Availability of Data and Materials

Data sharing is not applicable to this article as no datasets were generated or analyzed during the current study.

### Authors’ contributions

E.A.P and B.B. conceived and outlined this article. F.S. and B.B wrote this article. All authors approved this work.

## Acknowledgements

We would like to thank to Alexa Schmitz for critical reading of this manuscript.

